# The Prefrontal Cortex Organizes Visual Evidence by Representational Abstraction

**DOI:** 10.64898/2026.08.17.745311

**Authors:** Xinxu Shen, Jacob Miller, Ruedeerat Keerativittayayut, Chayanon Pamarapa, Chaipat Chunharas, Ioannis Pappas, Sirawaj Itthipuripat, Ian C. Ballard

## Abstract

Perceptual decisions often depend on both lower-level visual features and higher-level object representations, yet natural images entangle these levels of abstraction. We used deep neural network features to dissociate lower- and higher-level visual similarity during a naturalistic image-similarity judgment task. Participants relied on both forms of evidence, with greater weighting of higher-level similarity. Across prefrontal cortex, visual evidence was systematically organized by representational level: lower- and higher-level similarity were expressed in distinct prefrontal regions during comparison and remained spatially differentiated as they were transformed into decision signals. This organization was behaviorally relevant, with prefrontal activation predicting which level of evidence guided choice when the two conflicted. Together, these findings identify representational level as an organizing principle of prefrontal cortex during visual decisions.

## Introduction

Perceptual decisions are often described as choices based on sensory evidence (Gold & Shadlen, 2007). Yet in natural settings, the evidence available to a decision maker is rarely confined to a single level of representation. When deciding whether two faces belong to the same person or whether a car approaching in fog is familiar, decisions can be guided simultaneously by lower-level visual features and higher-level object or category representations. Although these sources of information can support the same choice, they can also diverge. A fundamental question is whether the brain preserves the distinction between these representational levels as visual properties are compared and evaluated during choice.

Although preserving multiple levels of visual representation may support flexible behavior, this organization remains poorly understood because many perceptual decisions can be resolved using a single dominant feature dimension, such as motion direction, appetitive value, or color. Additionally, lower- and higher-level visual features are often correlated in natural images—similar objects often appear visually similar—so behavioral and neural effects attributed to categorical or object-level information may also partly reflect lower-level image properties. Deep neural networks provide an opportunity to characterize and dissociate levels of visual similarity. Across successive layers, visually trained neural networks show a progression from local image structure to more abstract and object-level features. Layer-wise representations provide useful computational proxies for lower- and higher-level visual feature representations that correspond to functional organization in the visual system (Cichy et al., 2016; Khaligh-Razavi & Kriegeskorte, 2014; Yamins et al., 2014).

For visual information to guide choice, it must be represented in a form relevant to the current decision. For instance, in a similarity judgment, the relevant quantity is not what an image looks like in isolation but how it compares with another image. Lateral PFC maintains and selects task-relevant visual features, including location, shape, and object category (Christophel et al., 2018; Freedman et al., 2001; Lin & Lau, 2024; Miller et al., 1996; Rainer et al., 1998). It remains unclear whether PFC preserves the representational structure of individual visual inputs or reformats that information into task-relevant relationships between stimuli. Additionally, whether relational information is spatially organized according to the representational level of the visual information supporting it remains unknown. Prior work suggests that lateral PFC is organized according to task abstraction, with more posterior regions representing simpler feature-response relations and more anterior regions representing more abstract rules (Badre & D’Esposito, 2007; Badre & Nee, 2018; Koechlin et al., 2003; Miller & Cohen, 2001). However, sensitivity to lower- and higher-level visual similarity could vary across PFC even when the stimuli and task rule remain constant.

When making choices, relational visual information may be represented as evidence in favor of specific options. Midline PFC is broadly implicated in monitoring decision-relevant information and integrating signals that guide action (Rushworth et al., 2004; Shenhav et al., 2013), with ventromedial PFC (vmPFC) representing multiple decision-relevant inputs as a common subjective value signal (Bartra et al., 2013; Levy & Glimcher, 2012; Rangel et al., 2008). This common currency role may extend beyond explicit reward contexts to visual decisions, where the relevant variable is how strongly visual information favors each available option. Lower- and higher-level visual information may converge in the vmPFC as a common choice signal. Alternatively, choice signals reflecting visual information at different levels of representational abstraction may be spatially organized across mPFC.

We used deep neural network features (Simonyan & Zisserman, 2014) to design a stimulus set for a similarity judgment task that dissociates lower- and higher-level visual similarity (Ballard et al., 2026; Piriyajitakonkij et al., 2025). This design allowed us to investigate how different levels of visual representation influenced behavior and how they were organized during comparison and choice. Our results indicate that people use both lower- and higher-level information to guide visual similarity decisions. Multiple levels of visual information remain available across PFC rather than converging onto a single abstract representation. Representational level may therefore provide an organizing principle through which perceptual detail and abstract object-level information remain simultaneously available to support flexible behavior.

## Results

### Lower- and Higher-Level Visual Properties Drive Similarity Judgments

Participants performed a visual similarity judgment task in which they viewed one base image followed by two comparison images and selected which comparison image was more similar to the base image (Figure 1c). To dissociate different levels of visual information that may guide choice, we designed a stimulus set by parametrically varying visual similarity according to features extracted from different layers of the VGG16 deep neural network (Figure 1a; Simonyan & Zisserman, 2014; see Methods). This resulting stimulus set showed little correlation across trials between lower- and higher-level visual similarity (r = .052, Supplement Figure b). For example, one comparison image could be visually similar to the base image while belonging to a different category, whereas the other comparison image could share the same category but differ in visual appearance (Figure 1b). However, it is important to note that “lower”- and “higher”-level refer to relative positions within the VGG16 representational hierarchy, rather than to a strict distinction between early visual and semantic information. Most trials were comprised of stimuli from distinctly different categories and with different colors, shapes, and textures (Supplement Figure a). We then estimated the independent contribution of each representational level to similarity judgments.

**Figure 1.**
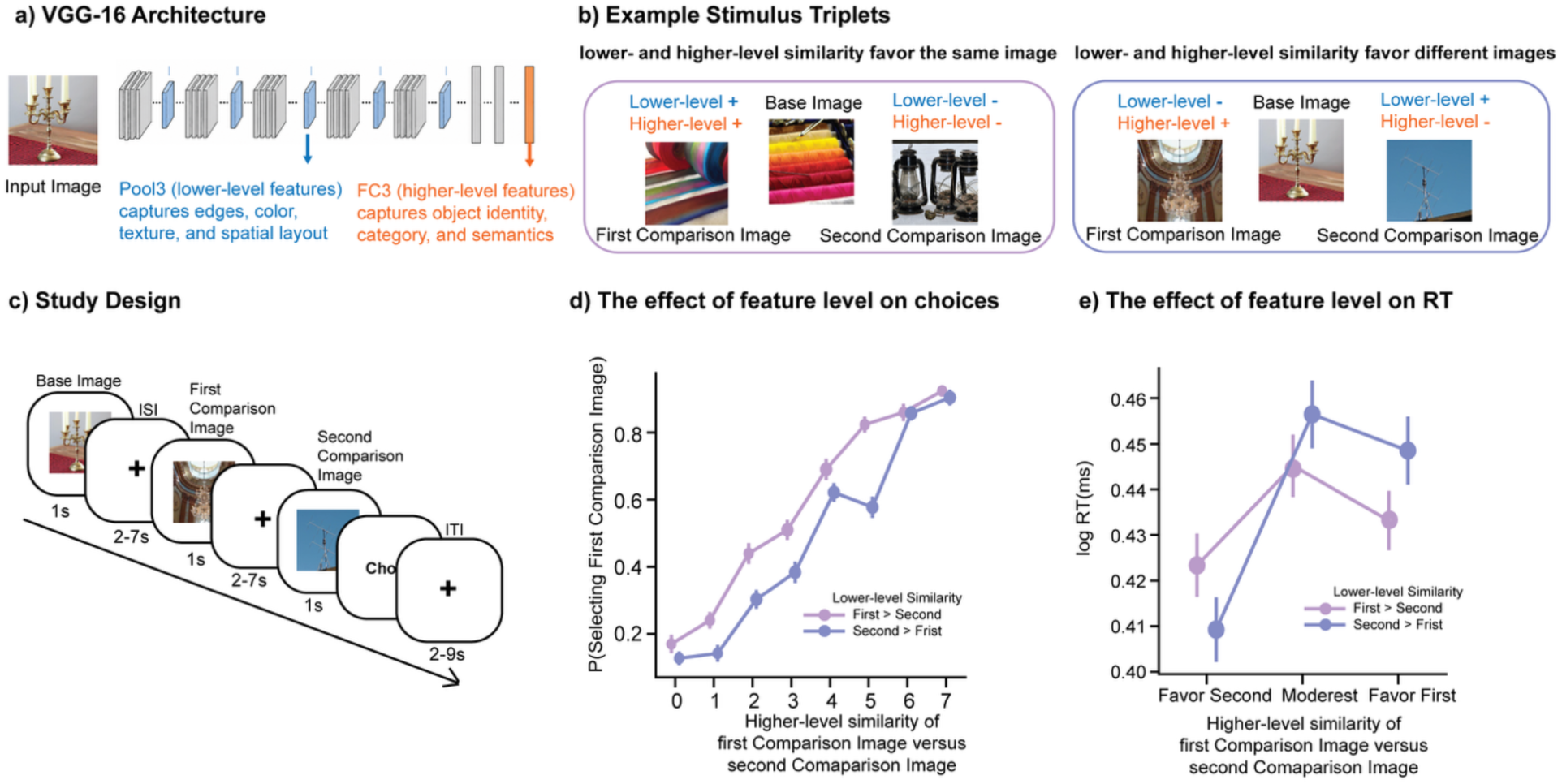
Study paradigm and behavioral dissociation of lower- and higher-level visual similarity. **a)** VGG16 feature hierarchy used to quantify visual similarity. Lower-level similarity was derived from Pool3 representations, which primarily capture lower-level visual features such as edges, color, texture, and spatial layout. Higher-level similarity was derived from FC3 representations, which primarily capture higher-level visual features, such as categorical information. **b)** Example stimulus triplets. Left, a trial example in which lower-level and higher-level similarity both favored the first comparison image. Right, a trial example in which lower-level and higher-level similarity favored different comparison images. **c)** Visual similarity judgment task. Participants viewed a base image, followed by two comparison images presented sequentially. After the second comparison image, participants indicated which comparison image was more similar to the base image. Twelve runs of 25 trials were completed during fMRI scanning. **d)** Probability of selecting the first comparison image as a function of the difference in higher-level similarity between the first comparison image and the second comparison image, relative to the base image. Both lower- and higher-level visual similarity independently predicted perceptual choice. Purple: trials on which lower-level similarity favored the first comparison image. Blue: trials on which lower-level similarity favored the second comparison image. Across the range of higher-level similarity, participants were more likely to choose the first comparison image if lower-level similarity also favored it, indicating that lower-level similarity contributed to choice. e) Log reaction time (ms) as a function of higher-level similarity difference. Error bars indicate standard errors.

Both lower- and higher-level visual representations independently predicted choices, suggesting that participants used multiple levels of visual information when making similarity judgments (Figure 1d). Greater higher-level similarity between the base image and the first comparison image, relative to the second comparison image, was associated with a higher likelihood of choosing the first comparison image (*β* = 1.36, *SE* = 0.04, *z* = 36.92, *p* < 0.001). Lower-level visual similarity also predicted choice (*β* = 0.26, *SE* = 0.03, *z* = 8.10, *p* < 0.001), although its influence was substantially smaller. There was no interaction between the influence of lower- and higher-level similarity on choice (*β* = -0.03, *SE* = 0.51, *z* = -0.71, *p* = 0.48).

We next examined conflict trials, in which the two levels favored different comparison images, to test how participants resolved competing visual evidence. On conflict trials, in general, participants were more likely to choose the higher-level-preferred image (intercept: *β* = 0.892, *SE* = 0.050, *z* = 17.69, *p* < 0.001). The probability of choosing the higher-level-preferred image increased as higher-level similarity evidence became stronger (*β* = 0.666, *SE* = 0.044, *z* = 15.20, *p* < 0.001). In contrast, the strength of the competing lower-level similarity evidence did not reliably predict choice of the higher-level-preferred image (*β* = -0.005, *SE* = 0.042, *z* = -0.12, *p* = 0.903), nor did it interact with higher-level similarity (*β* = 0.039, *SE* = 0.048, *z* = 0.82, *p* = 0.415). Thus, although both lower- and higher-level visual similarity contributed to choices overall, behavior was dominated by higher-level visual similarity when the two levels of evidence conflicted.

Finally, we asked whether lower- and higher-level visual similarity influenced the speed of the similarity judgment (Figure 1e). Because reaction times were right-skewed, we analyzed log-transformed reaction times. Higher-level similarity significantly predicted response time (*β* = 0.033, *SE* = 0.007, *t* = 4.83, *p* < 0.001). Lower-level similarity did not show a reliable main effect on response time (*β* = 0.006, *SE* = 0.007, *t* = 0.86, *p* = 0.388). However, lower- and higher-level similarity significantly interacted (*β* = -0.015, *SE* = 0.007, *t* = -2.18, *p* = 0.030), indicating that responses were faster when the two levels of similarity favored the same comparison image than when they provided conflicting evidence.

### Lateral Prefrontal Cortex Exhibits a Spatial Organization of Visual Feature Abstraction

The behavioral results show that participants used multiple levels of visual similarity to make decisions, with higher-level evidence carrying the stronger influence. We next asked how these levels of evidence were represented in PFC during image comparison. We first examined coding of image similarity during the presentation of the first comparison image, when participants had to maintain the base image and evaluate the relationship to the first comparison image. Prior work implicates the middle frontal gyrus (MFG) in maintaining and routing task-relevant information during working-memory and decision processes (Christophel et al., 2017; D’Esposito & Postle, 2015; Donahue & Lee, 2015). Given that both lower- and higher-level visual information influenced behavior, we also asked whether these levels of visual abstraction were represented in common or spatially distinct MFG regions.

Whole-brain analysis at the first comparison image revealed a spatial dissociation between lower- and higher-level visual information in MFG (Figure 2a). Greater higher-level similarity between the base and first comparison image was associated with increased activation in anterior MFG (*p* < .05, whole-brain FWE-corrected), whereas greater lower-level similarity was associated with increased activation in posterior MFG (*p* < .05, whole-brain FWE-corrected; see Supplementary Table 1&2 for additional regions).

**Figure 2.**
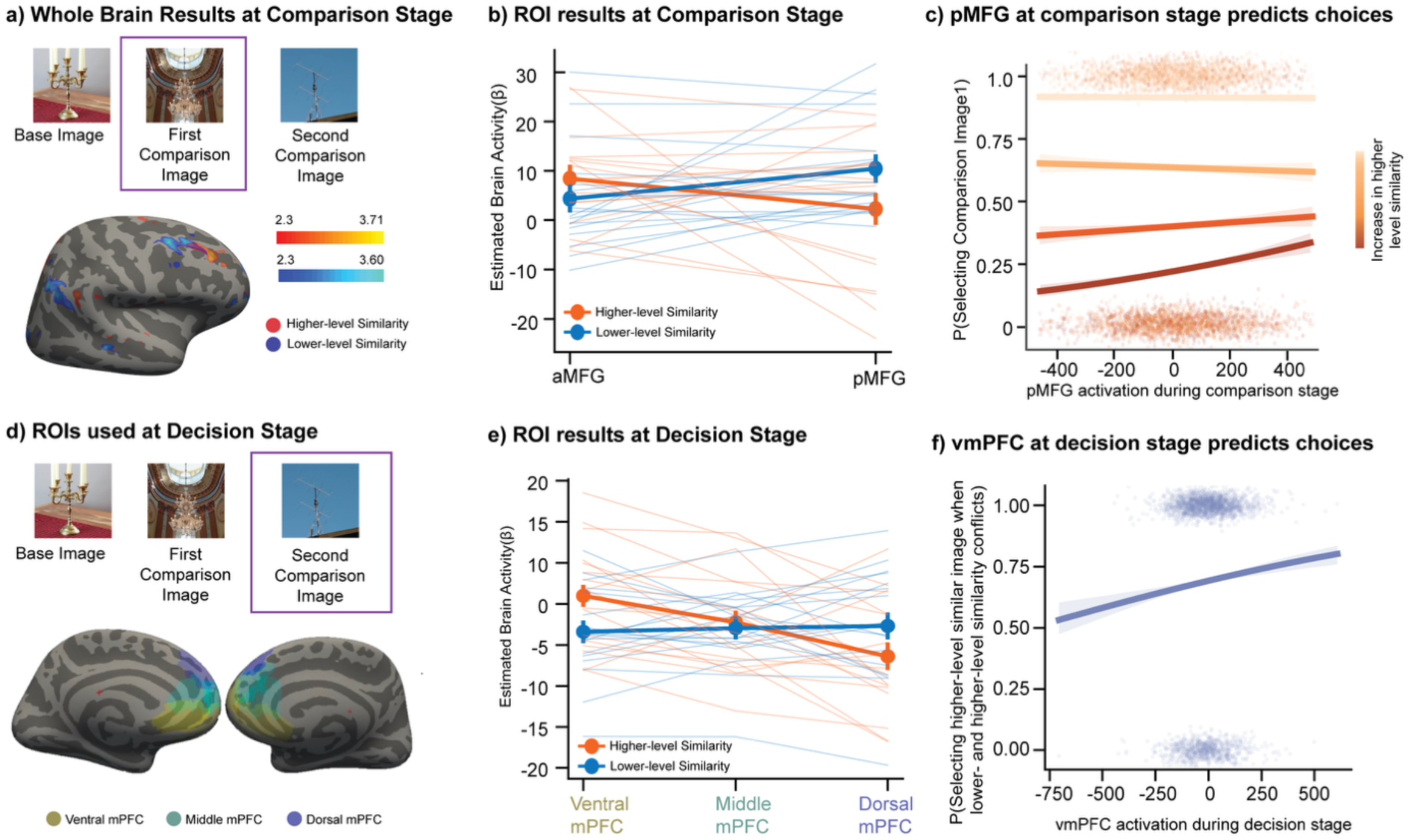
Segregated representation of relational visual information in lateral and medial prefrontal cortex. **a)** Whole-brain results during presentation of the first comparison image. Whole-brain activation maps showing regions modulated by higher-level similarity (red) and lower-level similarity (blue) between the base image and the first comparison image (Z > 3.1, cluster-corrected P < 0.05). Higher-level similarity selectively activated anterior MFG, while lower-level similarity selectively activated posterior MFG. **b)** ROI analysis of the lateral prefrontal cortex. Mean parameter estimates from anatomically defined aMFG and pMFG regions of interest revealed an interaction whereby lower- and higher-level visual information were represented more strongly in posterior and anterior MFG, respectively. Error bars indicate SEM. **c)** At the first comparison image, pMFG activity interacted with higher-level similarity in predicting choice. When higher-level similarity was low (dark orange), pMFG activity significantly predicted choice; when higher-level similarity was high (light orange), pMFG provided little additional predictive power. **d)** Medial prefrontal cortex ROIs used for decision-stage analyses. Ventral, middle, and dorsal mPFC regions were defined from the Schaefer atlas and grouped according to anatomical position along the ventral-dorsal axis. **e)** ROI analysis of medial prefrontal cortex during the decision stage. Mean parameter estimates for lower-level and higher-level similarity difference regressors, computed as the similarity of the chosen image minus the unchosen image, relative to the base image. Ventral mPFC preferentially tracked higher-level similarity and dorsal mPFC preferentially tracked lower-level similarity. Thin lines indicate individual participants; thick lines indicate group means. Error bars indicate standard errors. **f)** Trial-wise vmPFC activity on conflict trials predicted choice of the option favored by higher-level similarity, after controlling for the magnitude of the higher-level similarity advantage.

To further characterize this spatial dissociation, we analyzed responses in independently defined aMFG and pMFG regions of interest (ROIs; Figure 2b). A mixed-effects model with similarity level (lower-level vs. higher-level), ROI (aMFG vs. pMFG), and their interaction revealed a Similarity Level x ROI interaction (*β* = 12.26, *SE* = 5.05, *z* = 2.43, *p* = 0.015; see Supplementary Table 4 for main effects), indicating that aMFG and pMFG were differentially sensitive to lower- versus higher-level similarity. Pos-hoc tests indicated that pMFG responses were significantly greater for lower-level than higher-level similarity (*β* = 8.17, *SE* = 3.57, *z* = 2.29, *p* = .022), whereas aMFG showed numerically larger responses to higher-level than low- level similarity, though this difference did not reach significance (*β* = -4.09, *SE* = 3.57, *z* = -1.15, *p* = 0.252).

Together, these findings indicate that lower- and higher-level visual information are differentially represented across MFG, revealing a spatial organization of visual feature abstraction within lateral PFC. Importantly, the similarity signals identified in MFG reflected the relationship between the base image and the comparison image rather than the properties of either image alone. This raised a key question: did MFG simply represent features of current visual input, or did it recode visual inputs into comparison-relevant relationships?

### MFG encodes relational, not perceptual, similarity

To distinguish whether the MFG represented visual properties of the comparison image, akin to the ventral visual stream, or represented the task-relevant similarity between images, we performed a representational similarity analysis (RSA). As a validation step, we first examined whether the VGG-derived features captured established differences in representational abstraction across the ventral visual stream (Supplementary Figure c). Activation patterns in early visual cortical regions (V1-V3) reflected lower-level visual similarity between images, whereas higher-level visual similarity was weak or absent in these regions. Higher-order category-selective regions showed robust representation of higher-level similarity and weaker representation of lower-level similarity (Cichy et al., 2016; Guclu & van Gerven, 2015; Khaligh-Razavi & Kriegeskorte, 2014; Kriegeskorte et al., 2008). These findings replicate prior evidence that representations become increasingly abstract across the ventral visual stream and support the use of VGG-derived similarity measures to characterize neural activation patterns.

We next characterized perceptual and relational similarity representations in the ventral visual stream (Figure 3). Specifically, we examined whether image similarity patterns across trials were better explained by the perceptual similarity of comparison images or by the relational similarity between each comparison image and its base. In early visual cortex, neither relational nor perceptual similarity reached significance, although perceptual similarity showed a marginal effect (perceptual: *β* = .0053, *SE* = .0026, *t*(17) = 2.03, *p* = .059, 95% CI = [-.0002, .0109]; relational: *β* = .0039, *SE* = .0023, *t*(17) = 1.64, *p* = .119, 95% CI = [-.0011, .0088]). Direct comparison between relational and perceptual similarity revealed no significant difference (mean difference = -.0015, *SE* = .0019, *t*(17) = -.76, *p* = .458). In higher-order category-selective regions, both perceptual and relational similarity reached significance (perceptual: *β* = .0073, *SE* = .0029, *t*(17) = 2.52, *p* = .022, 95% CI = [.0012, .0134]; relational: *β* = .0115, *SE* = .0029, *t*(17) = 3.93, *p* = .001, 95% CI = [.0053, .0177]). Relational and perceptual similarity did not significantly differ from one another (mean difference = .0042, *SE* = .0034, *t*(17) = 1.26, *p* = .226). These findings suggest that activity patterns in higher-order category-selective regions reflect both image properties and task-relevant relationships between stimuli, consistent with evidence that the higher-order category-selective regions can encode relations among multiple objects as well as individual object properties (Abassi & Papeo, 2020, 2024).

**Figure 3.**
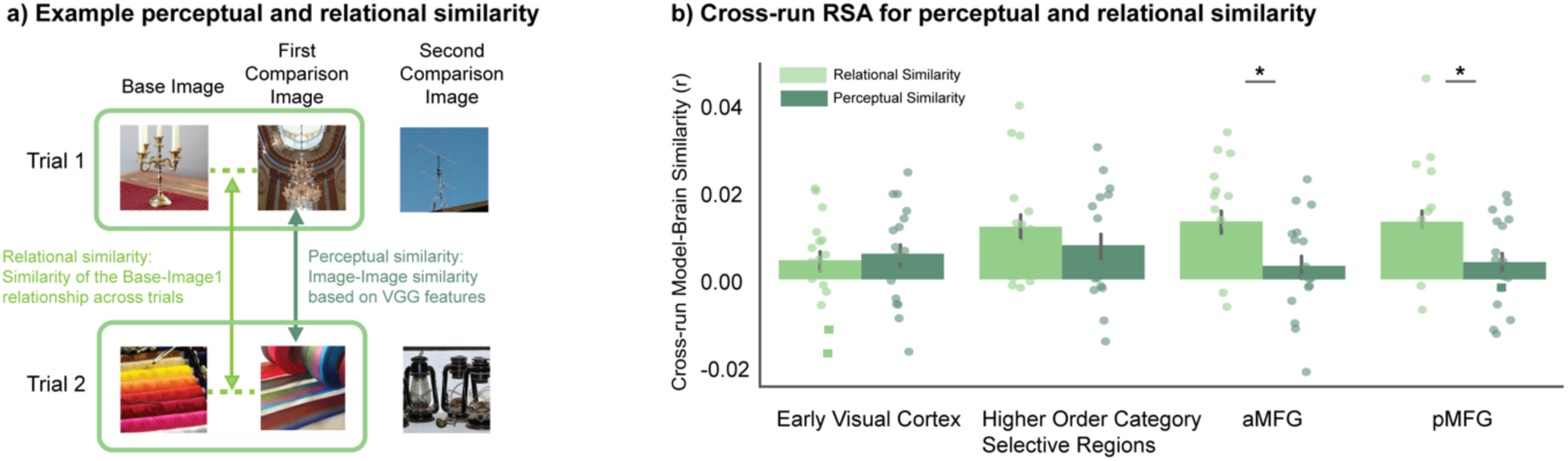
Middle Frontal Gyrus encodes relational, not perceptual, similarity. **a)** Two model representational dissimilarity matrices were constructed. The perceptual model quantified similarity among first comparison images across trials using VGG16 feature representations. The relational model quantified similarity in the base- first comparison image relationship across trials. Neural and model RDMs were compared using cross-run correlations only. **b)** Early visual cortex showed marginal perceptual coding and no relational coding. Higher-order category-selective regions encoded both perceptual and relational similarity. In contrast, both aMFG and pMFG selectively encoded relational similarity, with significantly stronger relational than perceptual coding. Error bars indicate standard errors. Asterisks denote significant within-region differences between relational and perceptual coding (p < 0.05).

**Figure 4.**
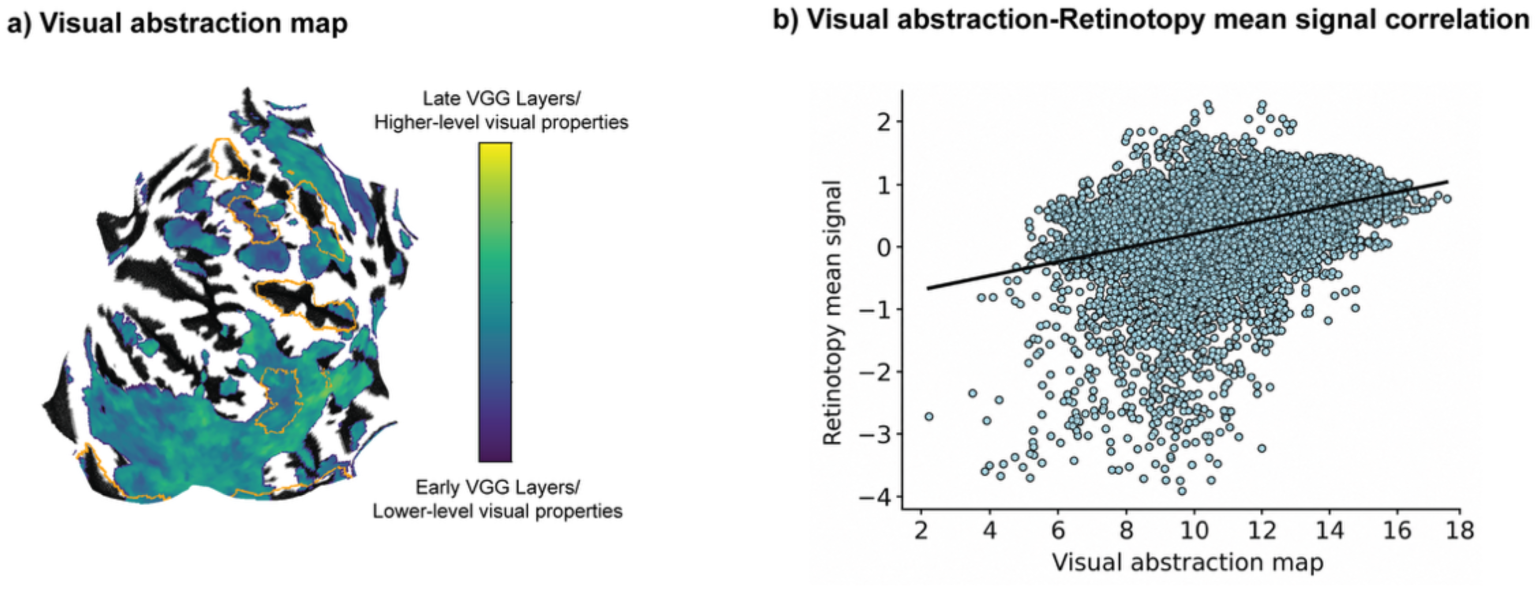
Correlation between visual abstraction and retinotopic organization. **a)** Cortical visual abstraction map showing the VGG layer that provided the best fit at each surface vertex. Cooler colors indicate earlier VGG layers associated with lower-level visual properties, whereas warmer colors indicate later VGG layers associated with higher-level visual properties. Orange contours indicate selected major sulci, including the superior and inferior frontal, central, intraparietal, fronto-polar and calcarine sulci. **b)** Significant correlation between the visual abstraction map and the retinotopy mean-signal map across cortical surface vertices. Each point represents one surface vertex, and the solid black line shows the fitted linear relationship.

In contrast, both anterior and posterior MFG preferentially represented relational rather than perceptual similarity. In aMFG, relational similarity was robustly significant (*β* = .0127, *SE* = .0028, *t*(17) = 4.61, *p* < .001, 95% CI = [0.0069, 0.0186]), whereas perceptual similarity was not (*β* = .0026, *SE* = .0026, *t*(17) = .98, *p* = .340, 95% CI = [-.0030, .0081]). Also, relational similarity was stronger than perceptual similarity within aMFG (mean difference = .01, *SE*= .0038, *t*(17) = 2.69, *p* = .016). A similar pattern was observed in pMFG: relational similarity was significant (*β* = .0127, *SE* = .0029, *t*(17) = 4.35, *p* < .001, 95% CI = [.0065, .0188]), perceptual similarity was not (*β* = .0034, *SE* = .0024, *t*(17) = 1.44, *p* = .167, 95% CI = [-.0016, .0084]), and relational coding was stronger than perceptual coding (mean difference = .0093, *SE* = .0036, *t*(17) = 2.56, *p* = .020). Thus, whereas the ventral visual stream contained both perceptual and relational information, MFG preferentially encoded comparison-relevant relationships between images rather than the perceptual properties of individual comparison images.

### Relational similarity signals in pMFG relate to choice

Having established that MFG carries relational information about lower- and higher-level similarity, we next asked whether these MFG signals were related to choice behavior. Theoretical accounts of PFC function suggest that representations are recruited when they are useful for behavior (Badre & D’Esposito, 2007). Accordingly, lower-level similarity signals in pMFG may be most predictive of choice when higher-level similarity does not clearly distinguish the options, whereas higher-level signals in aMFG may be most predictive when lower-level similarity is less informative. Consistent with the first prediction, mixed-effects logistic regression showed that both higher-level similarity (*β* = 3.20, *SE* = .09, *z* = 34.07, *p*< .001) and pMFG activity (*β* = .001, SE = 0.00, *z* = 4.014, *p* < .001) independently predicted subsequent choice. pMFG activity significantly interacted with higher-level similarity (*β* = -.001, *SE* =.001, *z* = -2.76, *p* = .006; Figure 2c): as higher-level similarity became more informative, the predictive influence of pMFG activity decreased. Thus, when higher-level similarity clearly favored one option, choices were primarily explained by the higher-level visual signal, and pMFG activity added little predictive value. In contrast, when higher-level similarity was weak or ambiguous, pMFG activity became a stronger predictor of choice. This pattern suggests that posterior MFG contributes lower-level relational information to decisions, particularly when higher-level evidence is insufficient to resolve the comparison. By contrast, the analogous interaction for aMFG was not significant (*β* = -.004, *SE* = .003, *z* = -1.07, *p* = .280), indicating that this flexible brain–behavior coupling was specific to posterior, rather than anterior, MFG

### Visual abstraction relates to known cortical organizational motifs

The MFG results suggest that relational visual evidence is spatially organized by abstraction level. We next asked whether this organization was specific to MFG or aligned with broader cortical visual organization. The Human Connectome Project 7T retinotopy dataset provides independent maps of several visual-field properties, including polar angle, eccentricity, and population receptive-field (pRF) size, as well as a mean-signal map reflecting visual responsiveness (Benson et al., 2018). We therefore constructed a whole-brain preferred visual abstraction map and compared it with the HCP retinotopy and mean-signal maps. For each voxel, we identified the VGG16 layer whose base-first comparison image similarity best explained comparison-stage BOLD responses. This yielded a cortical map in which lower values corresponded to lower-level visual features and higher values corresponded to higher-level features. Comparing the spatial organization of the visual abstraction map with HCP retinotopy maps allowed us to test whether the organization of relational abstraction mirrors known organizational motifs, such as pRF size, that may relate to the types of visual features represented at different levels of abstraction.

The visual abstraction map was not significantly correlated with polar angle, eccentricity, pRF size, or other classic retinotopic dimensions (all *p*s > 0.05), indicating that the organization of relational visual abstraction was not explained by basic visual-field organization. In contrast, the visual abstraction map was significantly correlated with the HCP mean-signal map (pearson r = 0.432, *p* = 0.002; spearman rho = 0.410, *p* = 0.002; null permutations via cortical spin-tests). Thus, the spatial organization of relational abstraction corresponded to broad variation in visual responsiveness but was distinct from the measured retinotopic dimensions. Together with the MFG findings, this suggests that relational visual abstraction is spatially organized in visually responsive cortex (Orban et al., 2004). This organization reflects variation in the level of image features associated with comparison-stage similarity signals rather than retinotopic position or receptive-field size. Additionally, exploratory analyses linking whole-brain functional-networks to this visual abstraction gradient are reported in the Supplement (Supplement Figure f).

### Decision-evidence abstraction varies along the mPFC dorsal–ventral axis

The prior analyses showed that lateral PFC organized relational visual information according to its level of abstraction during comparison. We next asked how these distinct forms of evidence were represented as participants compared the options and made a choice. The medial prefrontal cortex (mPFC) has been implicated in decision-making and in representing variables related to choice value and decision evidence (Bartra et al., 2013; Kable & Glimcher, 2007; Pisauro et al., 2017). Because vmPFC is often thought to integrate multiple inputs into a common value-like signal, one possibility is that lower- and higher-level visual evidence would converge in a shared vmPFC representation. Alternatively, the spatial segregation observed in MFG could persist into the decision stage, with dorsal and ventral mPFC differentially tracking lower- and higher-level decision evidence.

Whole-brain analysis during the decision stage revealed that vmPFC tracked higher-level decision evidence (Supplement Figure d), operationalized as the difference in higher-level similarity between the base image and the chosen versus unchosen stimulus (*p* < .05, whole-brain FWE-corrected; see Supplementary Table 3). No regions showed significant modulation by lower-level decision evidence. Our prior analyses revealed a spatial organization of relational visual information within MFG during comparison. Although the whole-brain analysis did not detect regions sensitive to lower-level visual decision evidence, we asked whether a similar spatial organization was detectable within mPFC during decision-making. We therefore conducted ROI analyses in independently defined ventral, middle, and dorsal mPFC ROIs (Figure 2d). A mixed-effects model revealed a significant Decision Evidence Level x ROI interaction (*β* = -8.08, *SE* = 2.44, *t* = -3.31, *p* = 0.001), indicating that the relative representation of lower- and higher-level decision evidence varied across mPFC (Figure 2e).

Within-ROI analyses revealed systematic variation in sensitivity to lower- and higher-level visual decision evidence across the dorsal–ventral axis of mPFC. In dorsal mPFC, activation varied more strongly with lower-level than higher-level visual decision evidence (*β* = 3.71, *SE* = 1.75, *t* = 2.12, *p* = .034). In middle mPFC, there was no significant difference in activation to lower- and higher-level decision evidence (*β* = −.71, *SE* = 1.73, *t* = −.41, *p* = .68). In ventral mPFC, the pattern reversed: activation varied more strongly with higher-level than lower-level decision evidence (*β* = −4.38, *SE* = 1.73, *t* = −2.53, *p* = .011), mirroring the whole-brain analysis. We additionally tested whether the regional differences varied linearly from ventral to dorsal mPFC by assigning the ventral, middle, and dorsal ROIs ordered position values. The interaction between decision-evidence level and ROIs was significant (*β* = 4.04, *SE* = 1.22, *z* = 3.32, *p* = .001), indicating that the relative sensitivity to higher- versus lower-level decision evidence varied systematically along the ventral–dorsal axis. Higher-level decision evidence was represented more strongly toward ventral mPFC, whereas lower-level decision evidence was represented more strongly toward dorsal mPFC.

The vmPFC finding raised a further question about how competing sources of decision evidence are adjudicated. If vmPFC simply reflects higher-level visual evidence, then its activity should covary with the objective strength of higher-level similarity. If vmPFC also contributes to the weighting of evidence during choice, then its activity should predict whether participants rely on higher-level similarity when lower- and higher-level evidence conflict. To test this second hypothesis, we focused on conflict trials in which lower-level and higher-level similarity favored different comparison images. Greater vmPFC activity during the decision period was associated with an increased likelihood of choosing the option favored by high-level similarity (*β* = 0.09, *SE* = 0.044, *z* = 2.042, *p* = .046), even after controlling for the magnitude of the high-level similarity (*β* = 2.05, *SE* = 0.14, *z* = 14.47, *p* < .001; Figure 2f). This result suggests that vmPFC activity reflected more than the objective strength of higher-level similarity evidence; it was also associated with a greater tendency to rely on abstract, object-level similarity when competing perceptual signals were present. These findings are consistent with a model in which the mPFC adjudicates how competing sources of visual information are weighted during decision making.

Together, the lateral and medial PFC findings suggest that representational level organizes visual information from comparison through choice. Along lateral PFC, relational visual evidence was organized from lower to higher levels of abstraction along the posterior– anterior axis. Along medial PFC, evidence at these different levels remained spatially differentiated as it contributed to choice. This organization allows both lower- and higher-level information to remain available to guide behavior.

## Discussion

In naturalistic settings, low-level features, object identity, and goals are often intertwined. Our findings suggest that prefrontal cortex does not rely on a single, fully abstracted representation of visual evidence. Instead, lower- and higher-level evidence remained concurrently available as spatially differentiated prefrontal signals during comparison and choice. Along lateral PFC, relational visual evidence was organized from posterior to anterior according to its level of abstraction during comparison, whereas along medial PFC, evidence at different levels was organized from ventral to dorsal in relation to choice. This organization provides a potential cortical architecture through which the same visual input can support decisions based on either perceptual detail or abstract object-level information, rather than being collapsed into a single representation.

Our finding of an anterior–posterior distinction in representational abstraction within MFG bears some resemblance to hierarchical control theories of lateral PFC, whereby posterior regions support concrete stimulus-response relationships and anterior regions support more abstract task rules or structure (Badre & D’Esposito, 2007; Badre & Nee, 2018; Koechlin et al., 2003; Ladwig et al., 2026). The present organization is distinct from that described in classical hierarchical-control paradigms, however, because the task rule—to judge which image was more similar to a reference image—remained constant across trials. What differed across prefrontal regions was the representational level of the visual evidence supporting the judgment, rather than the abstraction of the rule, context, or response policy. Thus, the present findings extend accounts of lateral PFC organization by showing that representational abstraction within the evidence itself can structure prefrontal activity under constant task demands.

However, our results do not exclude the possibility that aMFG and pMFG interact hierarchically during comparison, for example if higher-level similarity representations in aMFG modulate the processing or behavioral contribution of lower-level relational representations in pMFG. The finding that pMFG activity predicted choice most strongly when higher-level stimulus evidence was weak is compatible with lower-level relational information becoming more behaviorally relevant when object-level information is insufficient to resolve the comparison. Establishing an asymmetric or hierarchical interaction between aMFG and pMFG would require causal perturbation or analyses that directly assess interregional influence (Nee & D’Esposito, 2017).

MFG preferentially represented the relationship between each comparison image and the base image, indicating that visual information was organized according to its relevance for the current judgment. In contrast, higher-order visual cortex represented both image-centered and relational information. This relational format is important because perceptual choice depends not only on what each image looks like, but on how each image relates to the decision goal. In this sense, MFG appears to represent visual input in a goal-relevant, relational format. Notably, information about higher-level relational similarity was present in both aMFG and pMFG, consistent with the positive contribution of both lower- and higher-level evidence to univariate activation in the ROI analysis. Thus, the findings support an organization by dominant representational level rather than a strict regional dichotomy between lower- and higher-level relational visual information.

The cortex-wide abstraction analysis further suggested that organization by representational level was not isolated to MFG. A preferred-layer map indexing which level of the VGG hierarchy best accounted for comparison-related activation correlated with the HCP mean visual-response map, but not with eccentricity, pRF size, or the other tested retinotopic dimensions. Together, these findings suggest that abstraction-related similarity signals extend beyond MFG and align with broader visually responsive cortex.

During the decision phase, midline PFC showed a ventral–dorsal organization by abstraction level. Dorsal mPFC was more sensitive to lower-level chosen-versus-unchosen similarity, whereas vmPFC was more sensitive to higher-level chosen-versus-unchosen similarity. This pattern complements the lateral MFG organization observed during comparison, suggesting that different levels of visual information remain spatially organized as they are expressed as choice-related evidence. It also mirrors the behavioral finding that both lower- and higher-level similarity influenced choice. In this sense, the midline PFC organization may reflect a decision system that preserves multiple sources of visual evidence rather than reflecting a spatially uniform scalar signal. Further, vmPFC activation was associated with higher-level- aligned choices on conflict trials even after controlling for the magnitude of the higher-level similarity advantage. Thus, vmPFC activation reflected trial-to-trial variation in whether higher-level evidence guided behavior. Together, these findings identify a ventral-dorsal organization of midline PFC, in which perceptual decision evidence is structured by the abstraction level of visual information.

The chosen-minus-unchosen similarity signal resembles relative chosen-value signals commonly observed in economic decision-making tasks (Bartra et al., 2013; Levy & Glimcher, 2012; Rangel et al., 2008), although the task contains no feedback, reward, or explicit preference. It is possible that subjects may attributed value to higher confidence choices, which are likely to occur when chosen-minus-unchosen similarity is larger (De Martino et al., 2013; Lebreton et al., 2015). Alternatively, vmPFC may represent whichever option-relevant quantity links properties of the environment to goal-directed selection, even outside of value-driven choice (Hayden & Niv, 2021). In this view, the vmPFC conveys the relevant quantity for translating sensory evidence into behavioral action selection.

These findings show that people draw on information at multiple levels of visual abstraction to guide perceptual decisions and suggest that the spatial differentiation of lower- and higher-level features in sensory cortex is paralleled by an organization of relational and choice-related signals across prefrontal cortex. Keeping information at multiple levels of abstraction may enable the brain to use the same sensory input in different ways as behavioral demands change. This spatial organization offers one possible means of preserving both fine perceptual detail and more abstract knowledge for perceptual decisions.

## Methods

### Sample

Twenty participants (age range: 21–35 years; mean age: 24.35±3.22 years; 9 male participants) were recruited for the study. All participants provided written informed consent. The study was approved by the Institutional Review Board of Chulabhorn Royal Academy, Thailand, and was conducted in accordance with the Declaration of Helsinki. Although the sample size was modest for an fMRI study, the study included dense within-subject sampling, with 300 trials per participant across 12 blocks, allowing for more stable trial-wise estimates underlying our analyses. Task-fMRI reliability depends on both participant number and the amount of individual-level data available (Nee, 2019). All participants were included in the behavioral analyses. For fMRI analyses, data quality was assessed using MRI Quality Control (v24.0.0.0) metrics. Runs were excluded if mean framewise displacement exceeded 1.5 times the interquartile range upper bound, or if temporal signal-to-noise ratio fell below 1.5 times the interquartile range lower bound. Two subjects and six runs from three participants were excluded due to poor data quality.

### Stimuli

The stimulus set consisted of 300 triplets of naturalistic images, each comprising a base image and two comparison images. Triplets were drawn from the stimulus set of Piriyajitakonkij et al. (2025), designed to dissociate lower- and higher-level visual properties in natural images. Visual similarity between images was quantified using feature representations from two layers of VGG-16, a deep convolutional neural network. Lower-level visual similarity was derived from the Pool3 layer, which encodes edges, textures, and local shape properties and has been shown to approximate representations in early visual cortex (e.g., V2) (Piriyajitakonkij et al., 2025). Higher-level similarity was derived from the FC3 layer, which encodes abstract, object-level information and approximates representations in high-level ventral visual areas such as inferior temporal (IT) cortex (Piriyajitakonkij et al., 2025). Throughout this manuscript, “lower-level” and “higher-level” similarity refer to similarity computed from Pool3 and FC3 representations, respectively.

Because objects from the same semantic category often share both higher-level meaning and lower-level visual features, the stimulus set was constructed to minimize correlations between lower- and higher-level similarity across triplets (*r* = .052). Triplets were selected so that the two comparison images varied parametrically in their similarity to the base image along lower-level and higher-level dimensions. This resulted in trials in which lower- and higher-level similarity favored the same comparison image, as well as trials in which they favored different images (Supplement Figure b). This design enabled independent assessment of the contributions of lower- and higher-level visual information to visual similarity judgments and their neural substrates.

### Task and Procedure

Each participant completed 12 runs of a visual similarity judgment task during fMRI scanning, each run comprising 25 trials (300 trials total). On each trial, a base image appeared for 1 s, followed by a jittered interstimulus interval (ISI; 2–7 s, mean = 4.52 s). The first comparison image was then displayed for 1 s, followed by a second jittered ISI (2–7 s, mean = 4.52 s), and then the second comparison image for 1 s. The response window opened only after the second comparison image offset. Participants were instructed to select the comparison image they judged to be more similar to the base image “In the second part of the trial you will see two images. You need to determine which of these two images is more similar to the first image”. Intertrial intervals ranged from 2 to 9 s (mean = 4.4 s).

### Behavioral Analysis

Behavioral data were analyzed using mixed-effects logistic regression in *R* (v4.3.1; lme4 v1.1.34). For the choice analysis, participants’ choices were modeled as a binary outcome (1 = first comparison image chosen; 0 = second comparison image chosen). For each trial, we compared lower-level and higher-level visual similarity differences between the base image and the two comparison images as

*Δsimilarity = similarity(base, first comparison image) − similarity(base, second comparison image),*

separately for lower- and higher-level feature representations. These values were entered as fixed effects. An interaction term tested whether the influence of one similarity dimension depended on the other. A random intercept for participant accounted for between-subject differences in overall choice bias. The model specification was:

*choice ∼ Δlower-level similarity × Δhigher-level similarity + (1 | participant).*

For the reaction time analysis, because response times were positively skewed, choice response times were log-transformed before analysis. The standardized lower-level and higher-level visual similarity differences described above were entered as fixed effects. An interaction term tested whether the relationship between one similarity dimension and response time depended on the other. A random intercept for participant accounted for between-subject differences in overall response speed. The model specification was:

*Log(response time) ∼ Δlower-level similarity × Δhigher-level similarity + (1 | participant).*

Finally, for the conflict trial analysis, Conflict trials were defined as trials in which the higher-level and lower-level similarity differences favored different comparison images. Participants’ choices were modeled as a binary outcome indicating whether they selected the image favored by the higher-level representation (1 = higher-level-preferred image chosen; 0 = lower-level-preferred image chosen). The strength of higher-level evidence was calculated as the absolute higher-level similarity difference between the two comparison images, and the strength of lower-level evidence was calculated as the absolute lower-level similarity difference. Both evidence-strength measures were standardized and entered as fixed effects. An interaction term tested whether the influence of higher-level evidence strength depended on the strength of the competing lower-level evidence. A random intercept for participant accounted for between-subject differences in the overall tendency to choose the higher-level-preferred image. The model specification was:

*Choice_pref ∼ abs(Δlower-level similarity) × abs(Δhigher-level similarity) + (1 | participant).*

### MRI Data Acquisition

Participants underwent MRI scanning on a 3.0-T Ingenia MRI scanner equipped with a 32-channel head coil (Philips Medical Systems, Best, the Netherlands). Functional MRI data were acquired during the visual similarity judgment task using a standard gradient-echo echo-planar imaging sequence with the following parameters: repetition time (TR) = 2000 ms, echo time (TE) = 27 ms, flip angle = 90°, field of view (FOV) = 240 × 240 × 112 mm, acquisition matrix = 76 × 78, slice thickness = 4 mm, and voxel size = 3 × 3 × 4 mm. Task stimuli were presented via a mirror mounted inside the MRI scanner, and participants responded using a button box while viewing the stimuli. A high-resolution T1-weighted anatomical image was acquired using a sagittal 3D turbo field echo sequence with inversion preparation with the following parameters: TR = 8.1 ms, TE = 3.7 ms, flip angle = 8°, FOV = 240 × 240 × 200 mm, acquisition matrix = 240 × 240, voxel size = 1 × 1 × 1 mm, 200 slices, and no slice gap.

### MRI Preprocessing

Preprocessing was performed with fMRIPrep 23.2.1 (Esteban et al., 2019; RRID:SCR_016216), built on Nipype 1.8.6 (Gorgolewski et al., 2011; RRID:SCR_002502). Key steps are summarized here.

The T1w image was corrected for intensity non-uniformity with N4BiasFieldCorrection (ANTs 2.5.0; Tustison et al., 2010), skull-stripped using ANTs, and segmented into cortical and subcortical tissue classes with FSL FAST (Zhang et al., 2001). Spatial normalization to MNI152NLin2009cAsym space was performed using ANTs nonlinear registration with TemplateFlow (Ciric et al., 2022).

Functional data were motion-corrected with FSL mcflirt and coregistered to the T1w reference using boundary-based registration (6 degrees of freedom; Greve & Fischl, 2009). Confound time series included framewise displacement (FD), six head-motion parameters, whole-brain global signal, and the first six anatomical CompCor (aCompCor) components estimated by fMRIPrep; temporal derivatives and quadratic terms were included for motion and global signal regressors (Satterthwaite et al., 2013). Volumes exceeding FD > 0.5 mm or 1.5 SD DVARS were flagged as motion outliers. All resampling steps were combined into a single interpolation using cubic B-spline interpolation.

### Region-of-Interest Definition

Lateral PFC ROIs were derived from the Yeo 17-network functional atlas (Thomas Yeo et al., 2011): posterior MFG (pMFG) from parcel 97 and anterior MFG (aMFG) from parcel 83. Medial PFC ROIs were defined using the Schaefer atlas and grouped by anatomical position into ventral (parcels 168, 169, 379, 380), middle (parcels 173, 174, 381, 382), and dorsal (parcels 178, 385) subdivisions. Early and higher-order visual cortex ROIs were derived from probabilistic occipito-temporal visual cortex maps from Rosenke et al., 2021. The early visual cortex ROI consisted of areas such as V1d, V1v, V2d, V2v, V3d, V3v, whereas the higher-order category-selective regions consisted of regions associated with face, body, place, and character representations (mFus-faces, pFus-faces, IOG-faces, OTS-bodies, MTG-bodies, LOS-bodies, pOTS-characters, IOS-characters, CoS-places, TOS-places). All ROIs were defined in MNI space and resliced to each participant’s native functional space.

### General Linear Models

Run-level fMRI data were modeled using a fixed-effects general linear model (GLM) implemented in FSL. Event regressors modeled the onset of the base image, the first comparison image, and the second comparison image (duration: 1 s for the base and the first comparison image; 2 s for the second comparison image to include the decision period). Four parametric modulators were included to capture trial-by-trial similarity signals.

During the presentation of first comparison image, two parametric regressors captured similarity between the base image and the first comparison image, computed separately for lower-level and higher-level feature representations. During the presentation of the second comparison image and the decision phase, two parametric regressors modeled choice-related similarity signals. These were defined as the difference in similarity to the base image between the chosen and unchosen image:

*Δsimilarity = similarity(base, chosen) − similarity(base, unchosen),*

computed separately for lower-level and higher-level features. Parametric modulators were z-scored prior to model estimation. All regressors were convolved with a double-gamma hemodynamic response function.

Nuisance regressors included six head-motion parameters, framewise displacement, the first six aCompCor components, global signal, and cosine drift terms.

Within-participant models were estimated across runs using fixed-effects models. Group-level whole-brain analyses were performed using FLAME 1+2 mixed-effects modeling in FSL. Statistical maps were thresholded at Z > 3.1 with cluster-level FWE correction at *p* < .05.

### ROI Analyses

Mean parameter estimates for each similarity regressor were extracted from each ROI using *fslmeants* and averaged across voxels within each ROI. ROI-wise parameter estimates were submitted to group-level analyses in Python to test for differences in sensitivity to low- versus high-level visual similarity across regions.

### Layer-Specific GLMs and Gradient Mapping

To map the cortical gradient of visual feature abstraction, we constructed 21 additional first-level GLMs, one per VGG16 layer (conv1_1 through fc3, a total of 21 layers). Each GLM was identical to the primary GLM except that it included a single parametric regressor indexing base–first comparison image similarity computed from that layer’s feature representations. This approach allowed us to estimate, for each voxel, which level of the VGG-16 hierarchy best accounted for BOLD responses during the comparison stage.

For each voxel, we identified the layer yielding the maximum similarity beta estimate across the 21 GLMs and assigned the corresponding layer index (1–21) as a measure of preferred visual feature abstraction level. The resulting whole-brain visual abstraction gradient map index preferred visual abstraction level, where lower values indicate sensitivity to lower-level perceptual features and higher values indicate sensitivity to higher-level visual representations.

We then used the group preferred visual abstraction gradient map to characterize representational organization within the whole brain and to test whether it aligned with independent cortical reference maps. The visual abstraction gradient map was averaged across subjects, projected to the cortical surface, and correlated with the mean signal from the Human Connectome Project 7T retinotopic map (Benson et al., 2018). We used a cortical spin-test framework to test the correlation of these whole-brain maps against each other via the *neuromaps* analysis package (Markello et al., 2022). Additionally, we also calculated overlap with resting-state connectivity functional-network maps (Li et al., 2023), including the cingulo-opercular network.

### Single-Trial Analysis

For representational similarity analysis, we obtained single-trial beta estimates using the Least Squares-Single (LSS) approach (Mumford et al., 2012). A separate LSS model was constructed for each trial and image type (base image, first comparison image, second comparison image). LSS models were identical to the primary GLM except that: 1) a single regressor modeled the individual image of interest; 2) all remaining trials of the same image type were collapsed into a single nuisance regressor; and 3) no parametric regressors were included.

### Representational Similarity Analysis

For each participant and ROI, neural representational dissimilarity matrices (RDMs) were constructed from single-trial beta estimates. Single-trial beta estimates were demeaned across trials before similarity computation, to remove mean activation differences related to representational geometry. To reduce the influence of structured noise across voxels and temporal autocorrelation, demeaned single-trial beta estimates were pre-whitened using multivariate noise normalization (Walther et al., 2016). Pairwise neural dissimilarity was then calculated as 1 − Pearson correlation between multivoxel activity patterns.

Two model RDMs were constructed to capture distinct ways in which visual similarity could be encoded in representational structure. The *perceptual model* captured similarity across the first comparison image, defined as 1 − Pearson correlation between their higher-level VGG16 feature vectors: 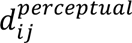 = 1 – *sim(first comparison image_i_, first comparison image_j_*. The *relational model* captured trial-wise differences in base–first comparison image higher-level similarity: for each pair of trials *i* and *j*, relational dissimilarity was calculated as the absolute difference in similarity value: 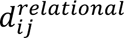 =∣ *S_i_* − *S_j_* ∣, where *s_i_*is the base–first comparison image similarity for trial *i*.

To reduce inflation from temporal autocorrelation, all RSA correlations were restricted to cross-run trial pairs. Neural and model RDMs were then vectorized, z-scored, and compared using linear regression. RSA was performed separately for each cross-run RDM pair within each subject and ROI, and the resulting beta coefficients were averaged across run pairs to obtain subject-level estimates for group analyses.

### Brain–Behavior Analysis

To test whether trial-wise prefrontal activity predicted perceptual decisions above and beyond objective image similarity, we conducted trial-wise brain–behavior analyses for both the first comparison image presentation period and the second comparison image decision period using Python. Single-trial parameter estimates were extracted from pMFG, aMFG, and vmPFC using the LSS models.

We used mixed-effects logistic regression to model binary choice as a function of neural activity, image similarity, and their interaction, with random intercepts for participants. Separate models were constructed for lower- and higher-level similarity. For the first comparison image presentation period, fixed effects included the lower- or higher-level similarity between base and the first comparison image, pMFG or aMFG activation (in separate models), and their interaction:

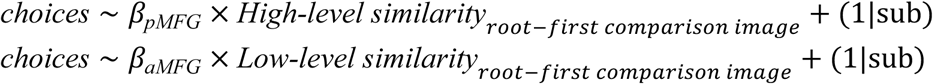

For the decision period, we focused on conflict trials, defined as trials on which the comparison image favored by lower-level similarity differed from the one favored by higher-level similarity. On these trials, we modeled whether participants chose the option favored by higher-level similarity as a function of vmPFC activity, controlling for the difference in higher-level similarity to the root image between the two comparison images:

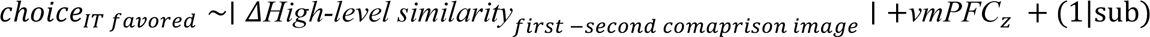

## Data and Code Availability Statement

The statistical analysis, figure generation code and behavioral data can all be found at https://github.com/KarenShen21/Deepgen_manuscript.git

## Artificial Intelligence Use Statement

Artificial intelligence assistance (ChatGPT, OpenAI; Claude, Anthropic) was used in revising the manuscript for grammar and clarity of exposition.

## Acknowledgements

We would like to thank the Ballard lab for helpful discussions regarding the results. This research has received funding support from by the National Research Council of Thailand (NRCT; fiscal years 2021–2024) to S.I., and C.C., Thailand Science Research and Innovation (TSRI; fiscal years 2021–2024: FRB670016/64, FRB660073/0164, FRB650048/0164, and FRB640008) to S.I., and from the NSRF via the Research and Innovation Acceleration Agency for Competitiveness and Area Development (RCAD;Program Management Unit for Frontier Brainpower and Future Industries; grant number B13F690044). We thank Kanyarat Benjasupawan and Praewpiraya Wiwatphonthana for their help with data collection and Maytus Piriyajitakonkij for technical assistance.

## Author Contributions

*Conceptualization*: ICB, IP, SI; *Data Curation*: XS; *Formal Analysis*: XS, ICB; *Funding Acquisition*: ICB, SI; *Investigation*: XS, RK, CP; *Methodology*: XS, IP and ICB; *Resources*: XS, ICB, SI; *Supervision*: ICB; *Validation*: XS and ICB; *Visualization*: XS, *Writing – Original Draft Preparation*: XS and ICB; *Writing – Review & Editing*: XS, JM, RK, CP, IP, SI and ICB. (CRediT).

## Supplement Information

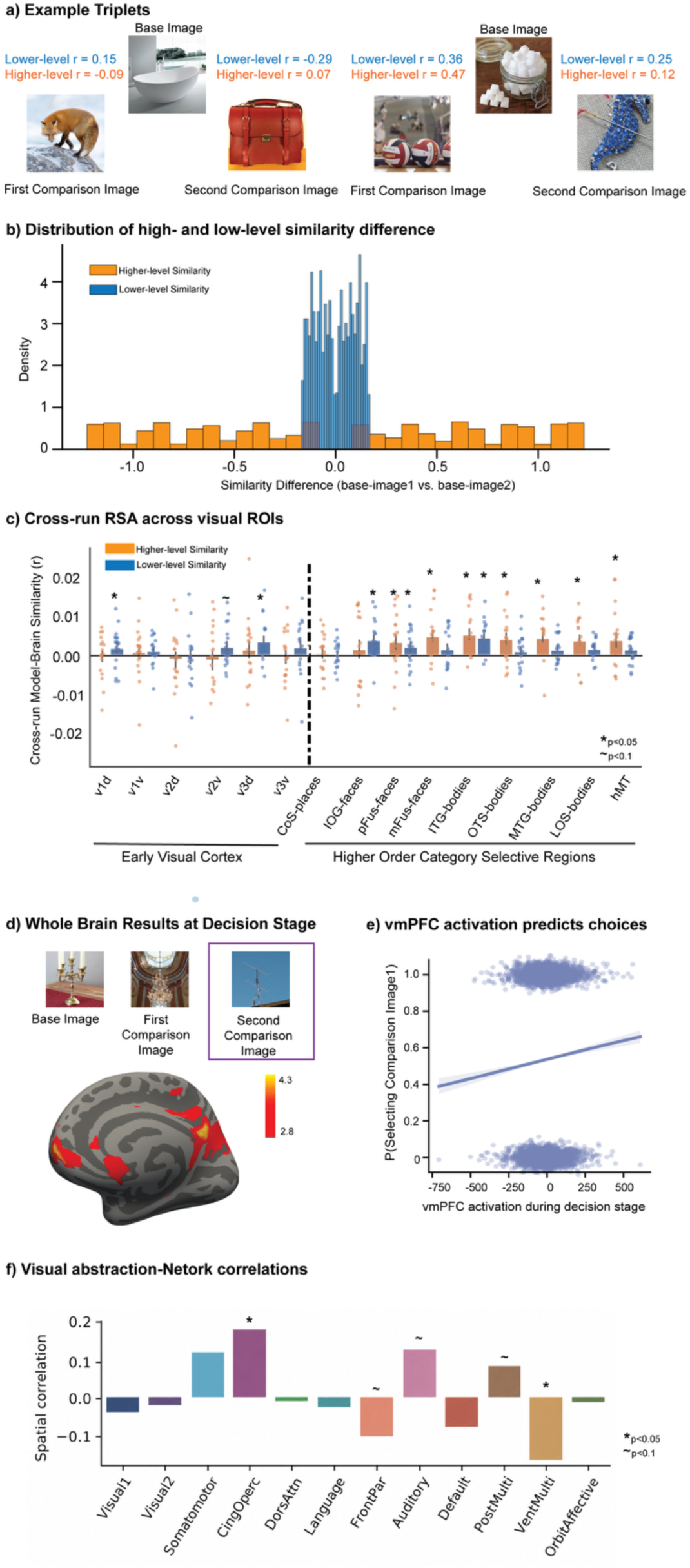

**a)** Example image triplets **b)** Distribution of higher-level and lower-level similarity differences across stimulus triplets. Similarity difference was computed as the similarity between the base image and the first comparison image minus the similarity between the base image and the second comparison image. Positive values indicate greater similarity for the first comparison image, whereas negative values indicate greater similarity for the second comparison image. Higher- and lower-level similarity differences were decorrelated across triplets, allowing their independent contributions to behavior and neural activity to be estimated. **c)** Cross-run representational similarity analysis across visual ROIs. Bars show group-average model–brain similarity between neural response patterns and model-derived similarity structure for higher-level and lower-level visual features. Dots show individual participants, and error bars indicate ± SEM. Cross-run RSA in visual ROIs from early visual cortex (V1–V3) to higher-order category-selective areas. Lower-level similarity was marginally represented in early visual regions, whereas higher-level similarity was reliably represented in higher-order category-selective regions, including face-, body-, and place-selective regions. **d)** Whole-brain decision-stage results. During the decision period, activity in vmPFC was associated with higher-level visual evidence for the chosen vs. unchosen image, relative to the base image. **e)** vmPFC activation during the decision stage predicted participants’ subsequent choices. Higher vmPFC activation was associated with a greater probability of selecting the first comparison image. **f)** Visual abstraction gradient is associated with large-scale functional networks. We tested whether the cortical visual-abstraction map, defined by the VGG layer that best explained comparison-stage activity at each surface location, was spatially related to independently defined functional networks from the Li et al. atlas. Bars show the correlation between the VGG- layer map and each functional network.

**Supplement Table 1.** Regions associated with higher-level similarity at comparison stage.

| Region | x | y | z | Z |
| --- | --- | --- | --- | --- |
| Anterior middle frontal gyrus | 31 | 33 | 30 | 3.67 |
| Precuneus | 7 | -64 | 66 | 4.01 |

x,y, and z are MNI coordinates, Z is peak z-score

**Supplement Table 2.** Regions associated with lower-level similarity at comparison stage.

| Region | x | y | z | Z |
| --- | --- | --- | --- | --- |
| Posterior middle frontal gyrus | 37 | 18 | 54 | 3.92 |
| Superior temporal gyrus | -64 | -26 | -2 | 3.84 |
| Orbitofrontal | 34 | 51 | -10 | 3.82 |

x,y, and z are MNI coordinates, Z is peak z-score

**Supplement Table 3.** Regions associated with higher-level similarity for chosen vs unchosen image at decision stage.

| Region | x | y | z | Z |
| --- | --- | --- | --- | --- |
| Cingulate cortex | 13 | -52 | 30 | 4.19 |
| Ventromedial prefrontal cortex | 4 | 65 | -6 | 4.33 |
| Precuneus | -11 | -64 | 30 | 3.99 |
| Lateral occipital cortex | 44 | -73 | 34 | 4.29 |
| Supramarginal gyrus | 63 | -26 | 38 | 4.12 |

x,y, and z are MNI coordinates, Z is peak z-score

**Supplement Table 4.** Mixed-effects model examining the effects of representational level and ROIs during comparison.

| Effect | $\beta$ | SE | z | p |
| --- | --- | --- | --- | --- |
| Intercept | 8.42 | 2.57 | 3.28 | 0.001 |
| Similarity level (Low-level vs. High-level) | -4.09 | 3.57 | -1.15 | 0.252 |
| ROI (pMFG vs. aMFG) | -6.16 | 3.57 | -1.73 | 0.084 |
| Similarity Level × ROI | 12.26 | 5.05 | 2.43 | 0.015 |

